# How to Extend the Capabilities of a Commercial Two-Photon Microscope to Perform Super-Resolution Imaging, Wavelength Mixing and Label-Free Microscopy

**DOI:** 10.1101/2021.09.02.458727

**Authors:** Chiara Peres, Chiara Nardin, Guang Yang, Fabio Mammano

## Abstract

Multimodal microscopy combines multiple non-linear techniques that take advantage of different optical processes to generate contrast and increase the amount of information that can be obtained from biological samples. However, the most advanced optical architectures are typically custom-made and require complex alignment procedures, as well as daily maintenance by properly trained personnel for optimal performance. Here, we describe a hybrid system we constructed to overcome these disadvantages by modifying a commercial upright microscope. We show that our multimodal imaging platform can be used to seamlessly perform two-photon STED, wavelength mixing and label-free microscopy in both *ex vivo* and *in vivo* samples. The system is highly stable and endowed with remote alignment hardware that ensures simplified operability for non-expert users. This optical architecture is an important step forward towards a wider practical applicability of non-linear optics to bioimaging.

## 1. Introduction

Imaging methods based on non-linear optical processes offer a combination of high contrast and sub-cellular resolution even in strongly scattering biological tissues, making them suitable for both *in vivo* and *ex vivo* applications [1, 2]. To exploit non-linear imaging processes in thick biological samples, the use of pulsed near-infrared (IR) excitation light is commonplace, because it allows high tissue penetration with low phototoxicity due to reduced photo-absorption. In addition, non-linear processes are excited mainly in the perifocal zone, implying intrinsic optical sectioning and reduced photodamage [3]. Non-linear microscopy allows flexible and non-invasive analysis in living tissues close to physiological condition, with high spatiotemporal resolution [4]. Moreover, different non-linear optical processes require similar pulsed near IR sources [5], making feasible, although not straightforward, to implement multiple imaging modalities in the same microscopy platform.

Second and third harmonic generation (SHG, THG, respectively) and coherent anti-stokes Raman scattering (CARS) are among the most widely used label-free non-linear imaging modalities. Harmonic generation is a coherent scattering process with no photon absorption, in which two (or more) photons interact with the sample to generate a photon of exactly half (for SHG) or one third (for THG) of the incoming wavelength [2]. SHG is due to the absence of inversion symmetry in the sample. The signal is typically generated by non-centrosymmetric molecules spatially aligned so as to add contributions and generate contrast. SHG is a particular case of sum frequency generation (SFG) [6] and is mainly used to investigate the structure of skin and bone collagen fibers, microtubules and myosin in muscles [7, 8]. THG depends on the discontinuity of third-order non-linear susceptibility that is typical of interfaces, allowing label free imaging of, for example, cell membranes, vessels, lipids and nuclei [9, 10]. CARS is based on molecular vibrational spectroscopy and allows selective non-invasive imaging by detection of specific chemical bond types and provides molecular fingerprinting [11]. Over the past several years, CARS had increased its popularity as an imaging method in the biomedical field, exemplified by imaging of lipids in biological sample and drug and/or cancer cell identification [12].

The most widely used non-linear method for biological imaging is two-photon excitation (2PE) (laser scanning) fluorescence microscopy (FM) [13], which allowed tissue penetration up to 1.6-mm depth in the mouse cortex [14]. In 2PE, two photons interact with the sample within ^~^5 fs to promote the fluorescent molecule under investigation from the ground state to an excited state. The excited molecule then emits a photon along the normal fluorescence emission pathway [15]. It is also possible to mix two IR excitation wavelengths for two-color two-photon excitation fluorescence microscopy (2c-2PEFM) [16]. In 2c-2PEFM, two different pulsed femtosecond laser are carefully spatially-aligned and synchronized so that the 2PE process can be promoted by either one of the two sources as well as by their combination. This technique allows for rapid multicolor 2PE of up to 3 fluorophores with efficient and independent control of the intensity of each fluorescence signal [16].

The substantially increased tissue penetration reached by two- [14] and three-photon microscopy [17], compared to single-photon excitation, comes at the expenses of spatial resolution due to the diffraction limit. Indeed, the use of IR light to elicit fluorescence almost doubles the linear dimensions of the excitation point spread function (PSF) compared to confocal one-photon imaging [18]. In order to improve the resolution of 2PEFM, in 2009 Moneron and Hell superimposed a donut-shaped STimulated Emission Depletion (STED) beam on the 2PE excitation beam, thus generating the first 2PE-STED architecture [19]. This mixed modality was used initially for super-resolution imaging in cell monolayers to disclose, for example, fine structure of GFP-tagged fixed cells [20]. However, the full potential of 2PE-STED become appreciable only when imaging fine structures in thick turbid tissue, where the 2PE advantages can be exploited. Thus, 2PE-STED was used, for example, to image the morphology of dendritic spines [21, 22] and both synapses and glial cells in living brain slices [23], or to analyze quantitatively dendritic spine morphology in a living mouse brain [24]. Later, Coto Hernandez et al. improved 2PE-STED performance by using a CW-STED laser, time-resolved detectors, and the SPLIT principle to discard non super-resolved photons [25]. A 2PE-STED system based on electrically controlled optical components, called “advanced easySTED”, was built by Otomo et al. with a simpler and compact architecture and shown to be more stable than prior implementations [26]. However, neither Coto Hernandez et al. nor Otomo et al. used their systems for 2PE-STED imaging in thick biological samples. Moreover, current multimodal or 2PE-STED microscopes, such as those quoted above, were based on custom-made architectures that require complex alignment procedures and daily maintenance by properly trained personnel [27]. To our knowledge, commercial multimodal 2PE-STED microscopes are not available.

Here, to partially fill this gap, we report the implementation of a highly reliable and stable multimodal imaging platform derived from a commercial upright microscope to perform 2PEF, 2PE-STED, 2c-2PEFM, SFG and CARS in *ex vivo* and *in vivo* samples. To ensure day-to-day serviceableness even for non-expert users, we endowed the platform with a system for remote alignment of critical components. This optical architecture is an important step forward towards a user-friendly multimodal microscope that can widen the practical applicability of non-linear optics to microscopy users interested in addressing biological questions *in vivo*.

## 2. Methods

### 2.1 Optical architecture

The system described in this article (Figure 1A, 1B, 1C, 1D) was based on a custom-made Leica TCS SP8 upright multi-photon excitation (MPE) scanning laser microscope (Leica Microsystem, Wetzlar, Germany). This system was designed for *in vivo* imaging deep in tissue and was optimized to perform three-dimensional super-resolution microscopy by one-photon Stimulated Emission Depletion (1P-STED). The infrared (IR) laser source was a tunable femtosecond pulsed Titanium Sapphire (Ti:Sa) Chameleon Ultra II Laser (Coherent, Santa Clara, CA, USA) with output range from 680 nm to 1080 nm. To mitigate group delay dispersion (GDD), which severely limits the performance of any two-photon system, the Ti:Sa laser hosted a negative pre-chirping unit. The Ti:Sa laser was coupled to a MPX, Chameleon Compact Optical Parametric Oscillator (OPO) (Coherent, Santa Clara, CA, USA) to extend the range of excitation wavelengths. The OPO provided two beam outputs: Ti:Sa output, 680-1080 nm and OPO output, 1000-1340 nm. Each output was pulsed at 80 MHz, with the OPO train pulses delayed by 5.4 ns relative to Ti:Sa pulses. Each beam, after reflection off silver mirrors, was separately launched into an electro optical modulator (EOM) (Qioptic, Waltham, MA, USA) for fast beam shuttering in the MHz frequency range. Out of the EOM, the combination of a λ/2 plate mounted in a rotary motorized holder, followed by a polarizing beam splitter (PBS) was used to modulate the intensity of each beam. Collimation and diameter of the Ti:Sa beam were controlled by inserting a 1.2× beam expander (BE) formed by a pair of achromatic doublets with 60 mm and a 50 mm focal lengths, respectively (AC254-060-B and AC254-050-B-ML, Thorlabs Inc., New Jersey, USA). With this choice of focal lengths, the beam filled completely the back focal aperture of the water immersion objective (HC IRAPO L25×/1,00 W motCORR, Leica). To compensate for the 5.4 ns delay mentioned above, the Ti:Sa beam was directed towards an optical delay line, composed of a pair of silver mirrors mounted in a retro-reflector configuration on a high-precision motorized linear stage (8MTL1401-300-LEn1-100, STANDA, Vilnius, Lithuania). This device enabled computer-controlled sub-micrometric variations of the optical path length, i.e. temporal delay adjustments in the order of 5 fs. This level of precision was essential to perform (i) wavelength mixing and (ii) STED with fluorophores excited by either IR excitation source.

**Fig. 1.**
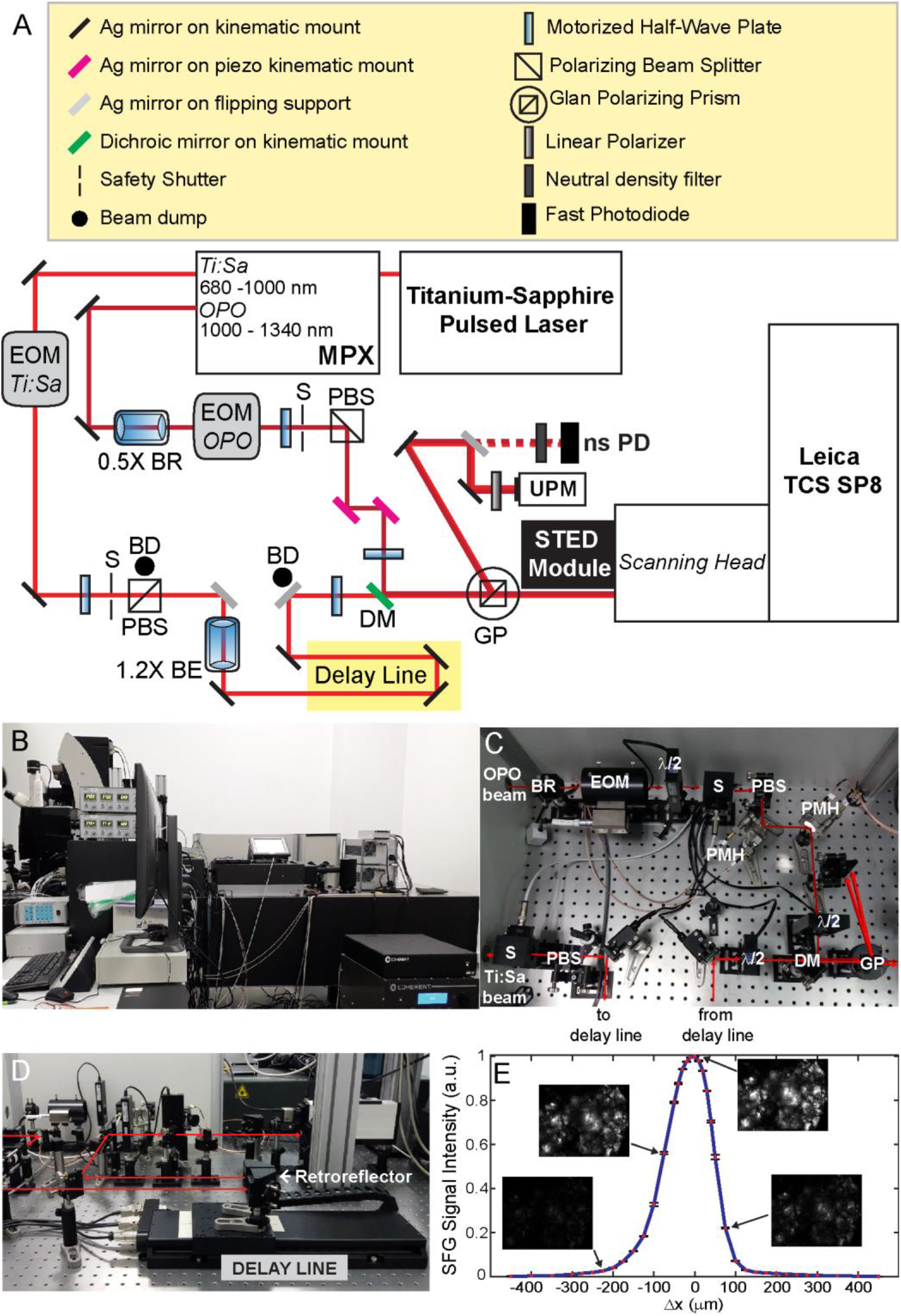
System description. **A**, Simplified optical scheme: Ag: Silver Mirror; EOM: Electro-Optic Modulator; PBS: Polarizing beam splitter; GP: Glan Prism; BE Beam Expander; BR: Beam Reducer; DM: Dichroic Mirror; S: Safety Shutter; BD: Beam Dump; UPM: Ultrafast Pulse Measurer; ns PD: nanosecond Photodiode; λ/2: Half-Wave Plate. **B**, Side view of the entire system. **C**, **D**, Detailed pictures of the beam routing components. **E**, Graph showing signal intensity of SFG signal excited by wavelength mixing in KDP crystals [16] vs. delay line length variation (Δx), where Δx=0 corresponds to the moving retroreflector assembly position at temporal synchronization. λ_1_ = 870 nm, P_1_ = 5 mW, λ_2_ = 1230 nm, P_2_ = 3mW. The SFG signal was detected in a narrow emission band between 505 and 515 nm. Insets are SFG images of the KDP crystals acquired at the indicated Δx values, keeping all the other acquisition parameters constant.

The OPO beam was stirred by using two remotely controlled piezo kinematic mirrors (POLARIS-K1S2P, Thorlabs Inc., New Jersey, USA), each coupled with an open loop piezo controller (MDT693B, Thorlabs Inc., New Jersey, USA). OPO beam stirring was essential for spatial alignment with the Ti:Sa beam after beam merging with a short pass dichroic mirror (SP1035, Leica). Before merging, each beam passed through a half wave-plate mounted on a remotely controlled rotary support. The merged beams impinged on a Glan polarizing prism (GPP, part number PGL 10.2, Bernhard Halle Nachfl. GmbH, Berlin, Germany) used to divert a fraction of the light towards a beam profiler (Grenouille 8-50-USB, Swamp Optics LLC, USA) and a high speed InGaAs photodetector (DET10N/M, Thorlabs Inc., New Jersey, USA). The latter was used for coarse synchronization of the two pulsed beams by feeding its output to a fast oscilloscope (part number HMO3004, Rohde & Schwarz GmbH & Co KG, Munich, Germany). The main transmitted component of each laser beam was aligned by means of a periscope in the Multi-Photon Port of the Scan Head of the upright Leica SP8 Microscope.

The commercial architecture was endowed with 3 different visible laser lines (488 nm, 532 nm and 635 nm) for one-photon multi-color imaging,4 internal descanned and 4 external non-descanned detectors. Each detector subsystem was composed of 2 InGasp photomultiplier tubes (PMT) and 2 higher sensitivity Hybrid Detectors (HyD). The microscope was endowed also with a commercial STED module (STED 3X, Leica Microsystem) for 1P-STED. The STED laser source was a 775 nm high power pulsed picoseconds laser source (Katana 08 HP, Onefive, Zurich, Switzerland). We modified the commercial system to trigger the STED laser with the electronics of the Ti:Sa laser with an intervening adjustable picosecond delayer.

### 2.2 System Alignment for two-photon STED

The commercial system allowed for automatic alignment of the STED beam and the 635 nm excitation laser. After running the automatic alignment procedure, we manually aligned the two IR pulsed laser beams with the 635 nm beam by inspecting the images of 1 μm multicolor fluorescent beads (T7282, TetraSpeck™ Microspheres, Thermo Fisher Scientific) diluted 1:10 in distilled water. To check the alignment of the IR pulsed beams with the donut shaped 775 nm STED beam, we used the reflection off 80 nm gold beads (see Section 3.1). For fine adjustment of the donut position and shape, we relayed on a customized feature of the control software that allowed us to move the vortex phase plate in two orthogonal directions within the STED module.

### 2.3 Animals

All animal experimentation was conducted in adherence to the NIH Guide for the Care and Use of Laboratory Animals and recommendations from both ARRIVE and PREPARE guidelines [28, 29]. Mice were bred and genotyped at Shanghai Biomodel Organism Science & Technology Development Co., Ltd., Shanghai (China). All experiments for this work were performed under animal production license sxck(Shanghai)2017-010 and animal usage license: sxck(Shanghai)2017-012. Mice were housed in individually ventilated caging systems at a temperature of 21 ± 2 °C, relative humidity of 55 ± 15% with 50–70 air changes per hour and under controlled (12 : 12 hour) light–dark cycles (7 am–7 pm). Mice had ad libitum access to water and a standard rodent diet. For most experiments, we used adult wild type C57BL6/N mice, both male and female, aged between 8 and 12 weeks. For some experiments we used also transgenic mice, both male and female, aged between 8 and 12 weeks, expressing the calcium (Ca2+) biosensor GCaMP6s in the Rosa26 locus after tamoxifen induction, generated by crossing the Jackson Laboratory strain #024106 (STOCK B6;129S6Gt(ROSA)26Sortm96(CAG-GCaMP6s)Hze/J) with the Cre-deleter strain #008463 (B6.129Gt(ROSA)26Sortm1(cre/ERT2)Tyj/J). For *in vivo* imaging, mice were anesthetized with 2% isoflurane and their temperature was kept at 37 °C by a thermostatically controlled heating pad. Animals were returned to their cages at the end of the imaging session.

### 2.4 Sample preparation

#### Tissue sample preparation

Mice were culled by trained personnel using gaseous anesthesia followed by a rising concentration of CO2 and cervical dislocation to confirm death, or cervical dislocation alone. For slice preparations of brain and kidney, each organ was explanted, quickly immersed in ice-cold PBS and cut in about 1 mm slices. Freshly excited slices were incubated for 30 min at 37°C in a solution of Nile Red (Cat. No. 72485-100MG, Sigma-Aldrich/Merck) pre-dissolved in methanol at a concentration of 1 mg/ml and diluted 1:20 in PBS. For skin specimen, we followed the protocol described in ref. [30]. Shortly, we excised a portion of dorsal skin comprising both epidermal and dermal layers (about 1 cm × 1 cm) and removed residual hair by using a hair removal cream. The organotypic skin culture was then maintained in a Trowell-type system, with the dermal side immersed in DMEM/F12 supplemented with 10% FBS and the stratum corneum exposed to air. The tissue was stained by applying a 400 μl drop of Nile Red 1 mg/ml diluted in methanol and incubated at 37°C for 30 min. The surface was then cleaned with ethanol and rinsed with PBS. After staining, samples were transferred under the microscope objective and imaged at room temperature for a maximum of 2 hours.

#### Immunostaining

Human HeLa cells plated on a glass coverslip were fixed with 4% formaldehyde in PBS for 15 min, washed 3 times with PBS and incubated for 30 min at room temperature with blocking buffer (3% bovine serum albumin and 0.1% Triton-X-100 in PBS). Cells were then incubated for 1 h at room temperature with the monoclonal mouse Anti-Tom20 (1:500, MABT166, Merck & Co., Kenilworth, NJ, USA). The antibody was revealed using Atto 594 goat anti-mouse IgG (1:500, 76085-1ML-F, Merck & Co., Kenilworth, NJ, USA). The coverslips were rinsed in PBS, mounted with prolong gold antifade medium (P36930, Thermo Fisher Scientific, Waltham, MA, USA) and sealed with nail polish.

### 2.5 Image acquisition and analysis

Throughout this article, λ denotes the wavelength of the laser beam propagating in empty space (unless otherwise stated). We define λ_1P_ and λ_2P_ as the excitation wavelength for 1PE and 2PE process, respectively. STED wavelength was fixed at 775 nm. P_1P_, P_2P_ and PSTED denote, respectively, the average 1P excitation, 2P excitation and STED laser power measured at the objective aperture. All images were acquired using the Leica Application Suite X Software. Fluorescence images were acquired with 63× glycerol immersion objective (HC PL APO CS2 63×/1.30GLYC) with 1.30 numerical aperture (NA). Unless otherwise stated, the pixel dwell time was 1.22 μs and each image was obtained by line averaging 32 times.

To acquire SFG, SHG, 2c-2PEFM, CARS and multimodal images, we used a 25× water immersion objective (HC IRAPO L 25×/1.0 W motCORR). We define λ_1_ and λ_2_ as the Ti:Sa and OPO excitation wavelength, respectively. P_1_ and P_2_ denote excitation power measured at the objective aperture for λ_1_ and λ_2_, respectively. For these imaging modalities, the pixel dwell time was 40.7 ns and each image was obtained by averaging 40 consecutive frames, unless otherwise stated.

Image processing was performed with the open-source software ImageJ/Fiji (ImageJ-win64). Matlab (R2019a, The MathWorks, Inc., Natick, MA, USA) was used for data extraction, analysis and fitting.

## 3. Results and discussion

### 3.1 Design, construction, spatial and temporal alignment of the multimodal optical architecture

To maximize the versatility of the commercial microscope adding multimodal capabilities, we implemented the system as shown in the scheme of Figure 1A. An overview of the system is presented in Figure 1B. The OPO provided two beam outputs: the Ti:Sa output (680-1080 nm) and the OPO output (1000-1340 nm). Each output beam was pulsed at 80 MHz, with the OPO train pulses delayed by 5.4 ns relative to Ti:Sa pulses. For spatial alignment of the two IR excitation sources, we rerouted the OPO beam by introducing two piezo kinematic mirrors in its optical path (Figure 1C), which allowed us to maintain spatial alignment via remote controls. To compensate for the 5.4 ns temporal delay, we inserted a motorized optical delay line in the Ti:Sa path (Figure 1D). To control beam divergence and to compensate for the extra length introduced by the delay line, we also inserted a 1.2× beam expander. The Ti:Sa and OPO beams merged at a 1035 nm dichroic mirror, followed by a Glan polarizing beam splitter used to divert a fraction of the light towards a beam profiler and a fast photodiode. For each beam, the fraction of diverted light was determined by the orientation of a motorized half-wave plate preceding the beam-merging dichroic mirror. By adjusting the delay line length while monitoring the fast photodiode output, we managed to synchronize coarsely the two IR pulsed beams at microscope input. Finer synchronization was achieved by focusing the objective on Potassium Dihydrogen Phosphate (KDP) crystals while further adjusting the delay line length to maximize the SFG signal elicited by the wavelength mixing phenomenon [16] (Figure 1E). The smoothness of the operation was guaranteed by sub-micrometric computer control of the motorized delay line.

A major goal for this project was to perform 2PE-STED. To this end, we triggered the 775 nm pulsed STED laser by means of the TTL output of pulsed Ti:Sa (after synchronizing the Ti:Sa and OPO pulses). A picosecond delayer was used to adjust the timing of the STED pulses relative to the Ti:Sa pulse.

To check the spatial alignment of Ti:Sa pulsed beam with the beam emerging from the 3D STED module, we used 80 nm gold beads plated on a microscope slide mounted on the Super Z Galvo stage. To obtain the *x-y* projection of the PSF, beads where imaged while performing standard raster scanning in the focal plane. To obtain a projection of the PSF in a plane that contained the optical axis, the stage was vibrated in the Z direction while performing fast line scans across the bead. The result of the alignment for both single and two-photon STED are displayed in Figure 2. Image quantification yielded the resolution values in Table 1.

**Fig. 2.**
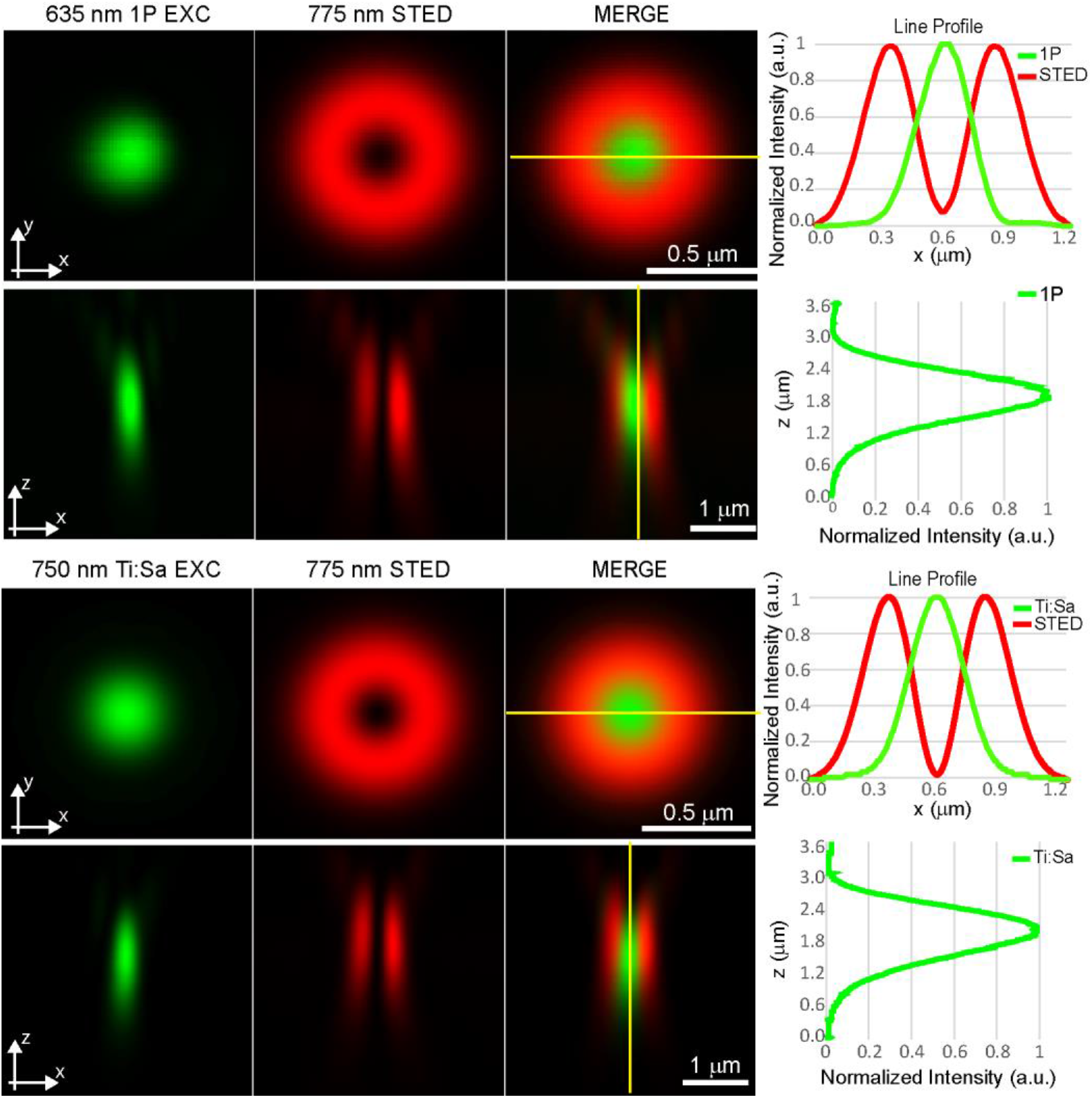
PSF characterization by means of beam reflection off of 80 nm gold beads. *x-y* and *x-z* projections of the circularly symmetrized excitation (green) and STED (red) beam PSFs for single photon beam at 635 nm (1P EXC, top) and Ti:Sa beam at 750 nm (Ti:Sa EXC, bottom). Graphs on the right are normalized intensity profiles measured along the yellow lines traced on the merge images.

**Table 1.** Full Width at half maximum (FWHM) of the measured PSFs in the *x-y* and *x-z* plane

| PSF | 1P EXC<br>(nm) | Ti:Sa EXC<br>(nm) |
| --- | --- | --- |
| x-y | 282 ± 2 | 311±6 |
| x-z | 889±9 | 1105±20 |

### 3.2 Characterization of the 2PE-STED performance of the system

To test the super-resolution performances of the temporally and spatially aligned system, we first used a fixed sample of HeLa cell mitochondria labelled with Tom 20 antibody counterstained with Atto 594 (Figure 3). With this type of specimen, clearly there was no advantage of using the 2PE-STED configuration compared to the ordinary 1PE-STED configuration.

**Fig. 3.**
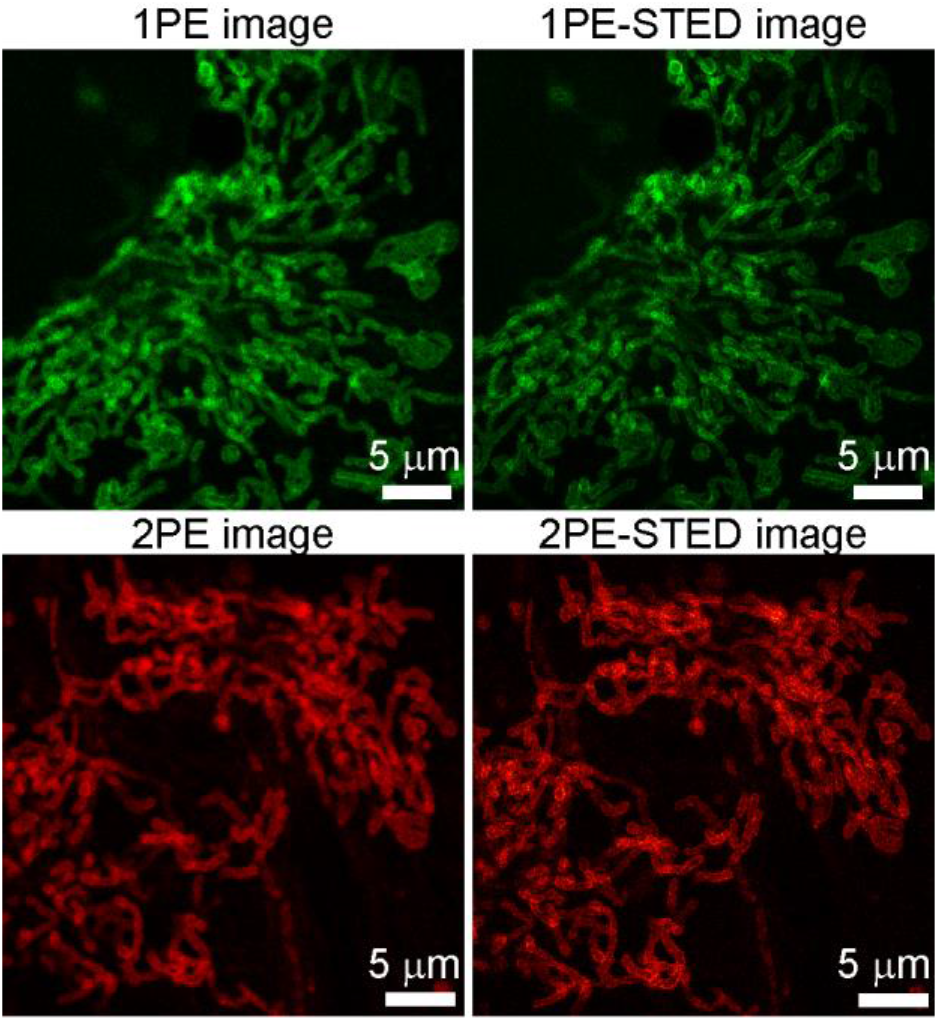
Comparison of one-photon excitation (1PE) and two-photon excitation (2PE) stimulated emission depletion (STED) images of fixed HeLa cell mitochondria stained with Atto 594. λ_1P_ = 532 nm, P1P = 10 μW; λ_2P_ = 850 nm, P2P = 15 mW; P_STED_ = 45 mW; fluorescence signal was acquired from 630 nm to 690 nm; pixel size = 45 nm; pixel dwell time = 2.4 μs.

However, this is not the case as soon as the sample thickness exceeded few tens of microns. In fact, by imaging myelinated structures in the white matter [31] of freshly dissected mouse brain slices, we observed that activating the STED beam in the 1PE-STED configuration caused loss of contrast, resulting in a blurred and poorly resolved image (Figure 4). Brain slices were stained with Nile Red, a fluorescent hydrophobic dye that quickly labels lipid structures and exhibits fluorescence only in a lipid environment [32]. Of note, Nile Red-stained beads were used to characterize the performance of the first 2PE-STED microscope [19].

**Fig. 4.**
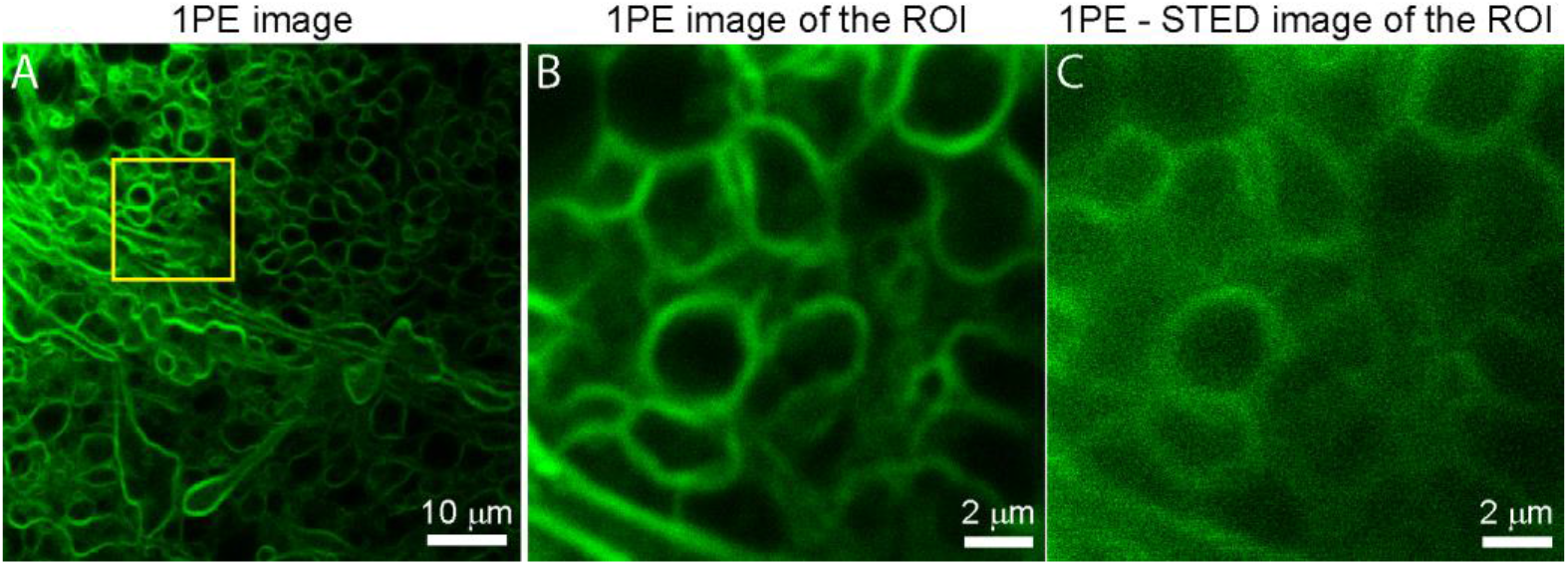
1PE and 1P-STED fluorescence images of myelinated structures in freshly excised mouse brain slice stained with Nile Red, acquired at 20 μm of depth in a wide collection band from 550 nm to 700 nm. λ_1P_ = 532 nm, P_1P_ = 15μW; P_STED_ = 42 mW; Pixel size: 90 nm (**A**) and 18 nm (**B**, **C**).

In contrast, activating the STED beam while performing 2PEFM on the same brain sample, at the same depth and at the same STED power, visibly improved both contrast and resolution (Figure 5).

**Fig. 5.**
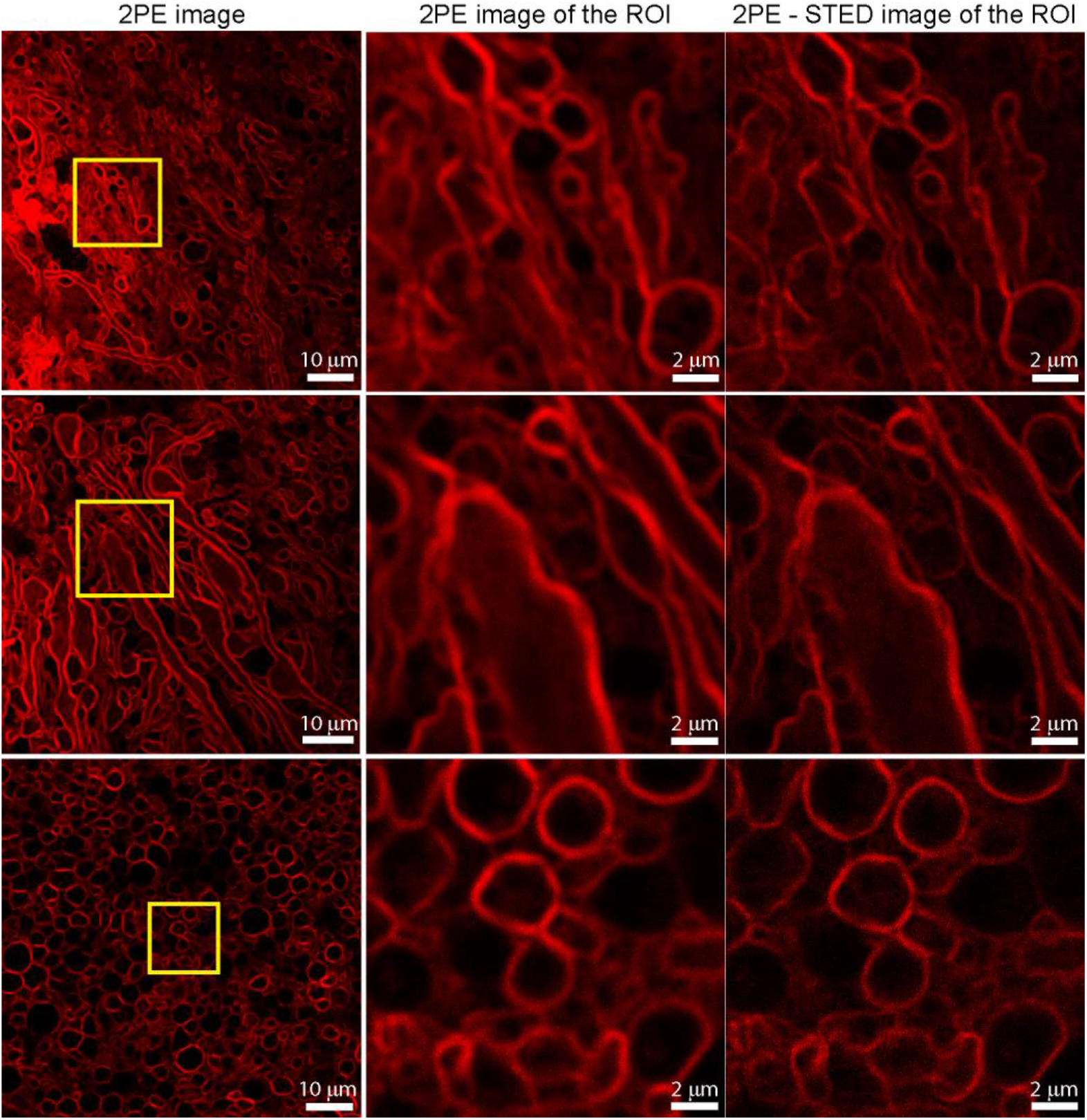
2PE and 2PE-STED fluorescence images of myelinated structures at 20 μm of depth in freshly excised white matter of mouse brain slices stained with Nile Red. λ_2P_ = 920 nm, P_2P_ = 20 mW; P_STED_ = 42 mW; fluorescence acquisition band = 550-700 nm; pixel size = 90 nm (first column) and 18 nm (all other images).

To evaluate the resolution improvement, we analysed the pixel intensity profile along selected lines intercepting fine structures of different freshly dissected samples (brain, kidney and epidermis) labelled with Nile Red (Figure 6; images acquired at depths comprised between 20 μm and 60 μm).

**Fig. 6.**
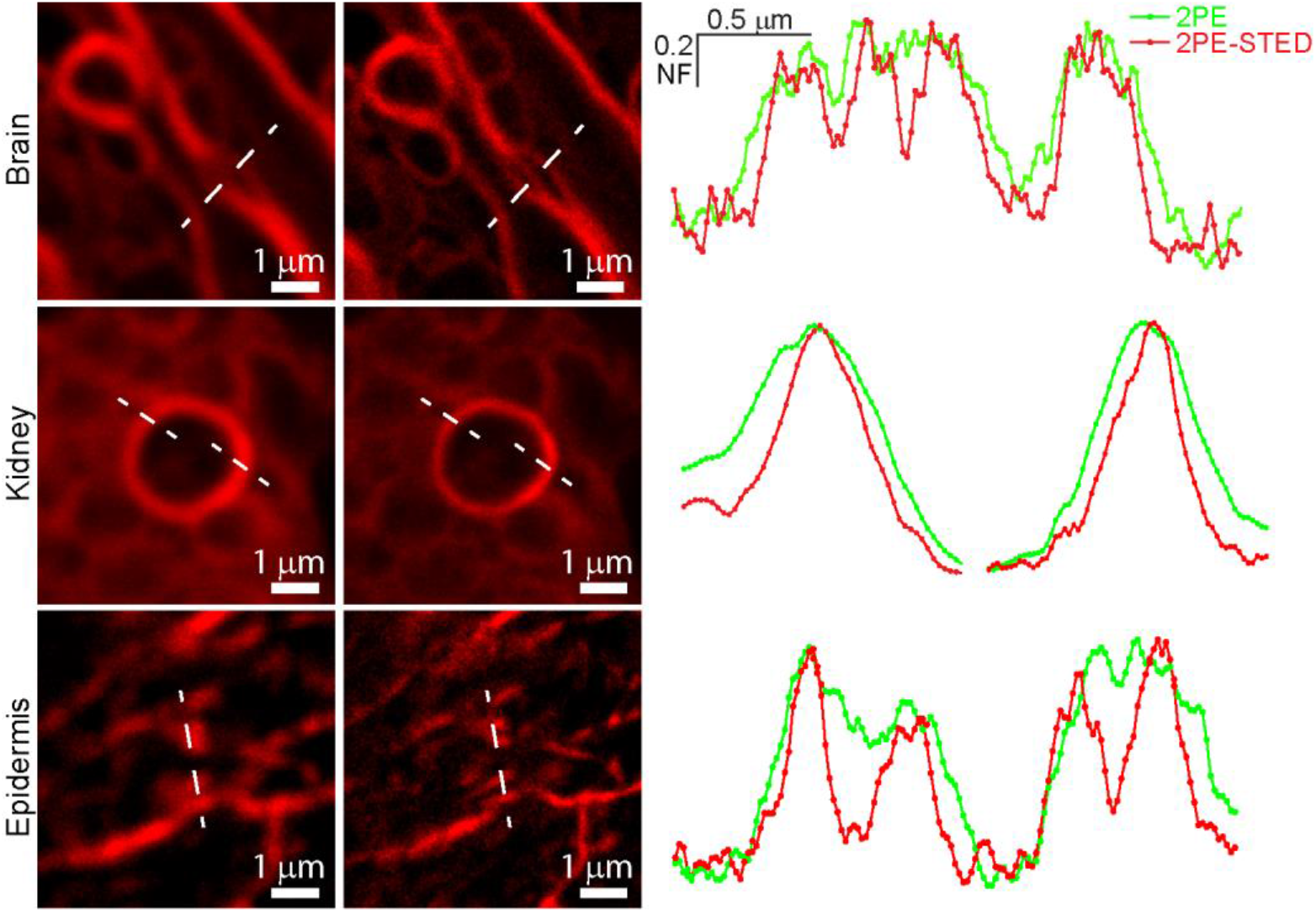
Resolution improvement due to 2PE-STED microscopy of fine structures in *ex vivo* tissues stained with Nile Red at depths comprised between 20 μm and 60 μm. Images on the left refer to 2PE and 2PE-STED of lipid structures in brain, kidney and epidermis samples. Graphs on the right are corresponding intensity profiles along the white dashed lines drown on the images at left, showing resolution improvement. λ_2P_ = 920 nm, P_2P_ = 23 mW; P_STED_ = 42 mW; fluorescence detection band: 550-700 nm; pixel size = 18 nm.

By combining 2PE-STED and SHG, we were able to image Nile Red-stained sebaceous glands nested in dermal collagen fibers both in freshly excited mouse skin and *in vivo* at a depth of 60 μm to 80 μm (Figure 7).

**Fig. 7.**
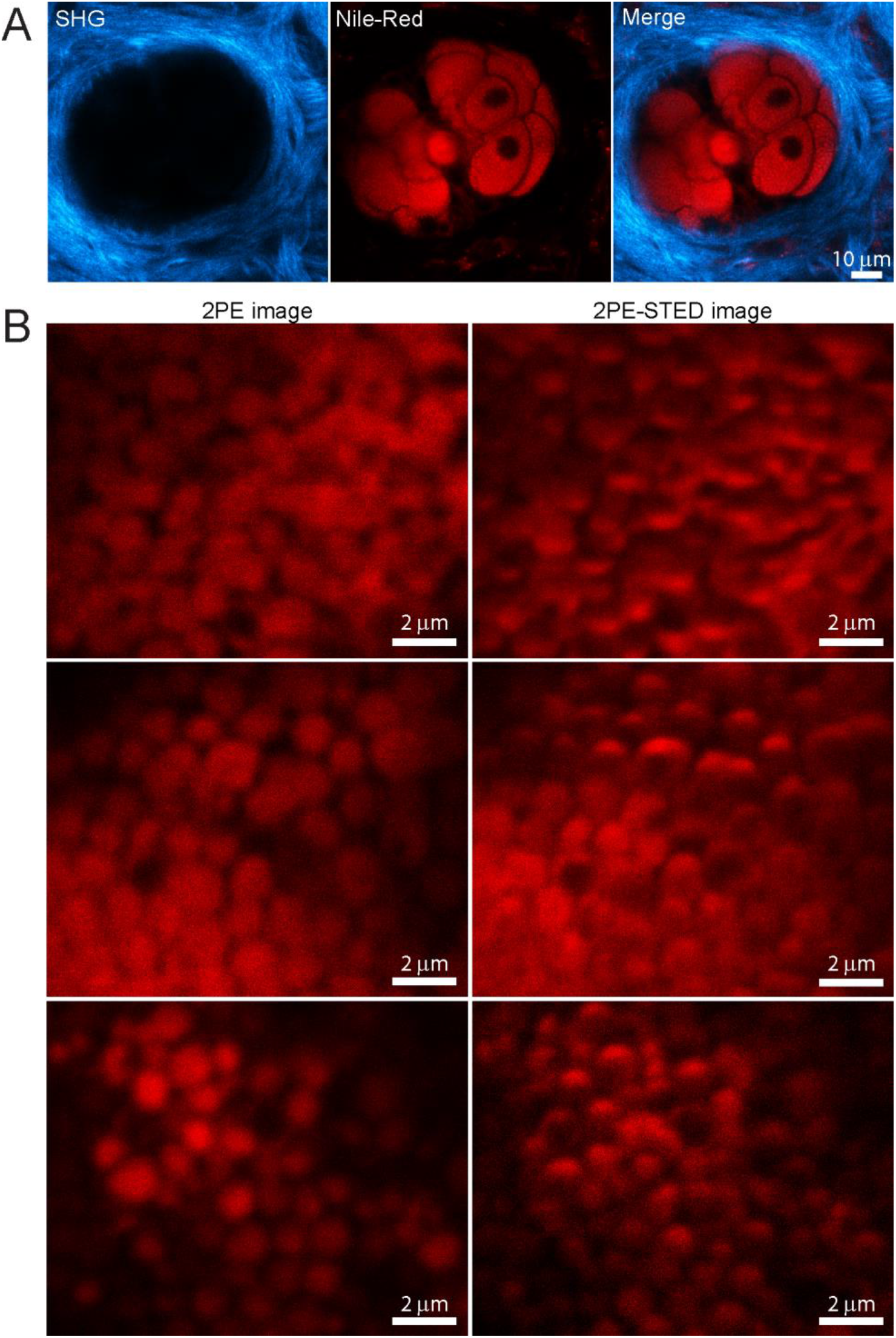
High-resolution imaging of sebaceous glands by 2PE-STED microscopy. **A**, Representative image of the *x-y* optical section of a Nile Red-stained sebaceous gland more than 70 μm deep in the earlobe skin of a live mouse. SHG signal from collagen fibers is shown in cyan. Images were acquired using a 25× water immersion objective at an excitation wavelength of 1080 nm; each image is the averages of 20 consecutive frames. P_2P_ = 10mW; pixel size =310 nm. Pixel dwell time = 1.367 ns. **B**, Comparison between 2PE and 2PE-STED images of sebocytes acquired 90 μm deep in mouse dorsal skin. Images were acquired with a pixel dwell time of 600 ns and averaging 32 consecutive frames. λ_2P_ = 920 nm, P_2P_ = 29 mW; P_STED_ = 52 mW; fluorescence detection band: 550-700 nm; pixel size = 18 nm.

### 3.3 Use of the system for *in vivo* label-free imaging of skin structures

To expand the full capabilities of the system over the 2PE-STED super-resolution imaging, we used additional optical components to exploit non-linear optical phenomena such as 2c-2PEM and CARS microscopy. In particular, the OPO not only extends the excitation wavelength range of the Ti:Sa laser, but can also be used to generate non-linear phenomena, based on the spatio-temporal synchronization of the two excitation laser beams (as mentioned in the Introduction).

Thus, we exploited the multimodal capabilities of our platform with the Ti:Sa laser tuned at λ_1_ = 840 nm and the OPO tuned at λ_2_ = 1105 nm to image the earlobe skin of live anesthetized mice ubiquitously expressing the green-fluorescent GCaMP6s indicator. With this wavelength combination, the system concurrently allowed us to (i) generate SHG signals from dermal collagen; (ii) visualize GCaMP6s fluorescence in keratinocytes and blood vessels by 2c-2PEFM, according to the equation 954 nm [16]; (iii) perform label-free imaging of sebocytes in sebaceous glands (Figure 8), as well as subdermal fat of adipose tissue (Figure 9) as a result of CARS. Note that, as mentioned in the Methods section, λ denotes the free-space wavelength whereas the corresponding wavelength in the tissue is unknown because we did not measure the refractive index of the tissue and its dispersion relations.

**Fig. 8.**
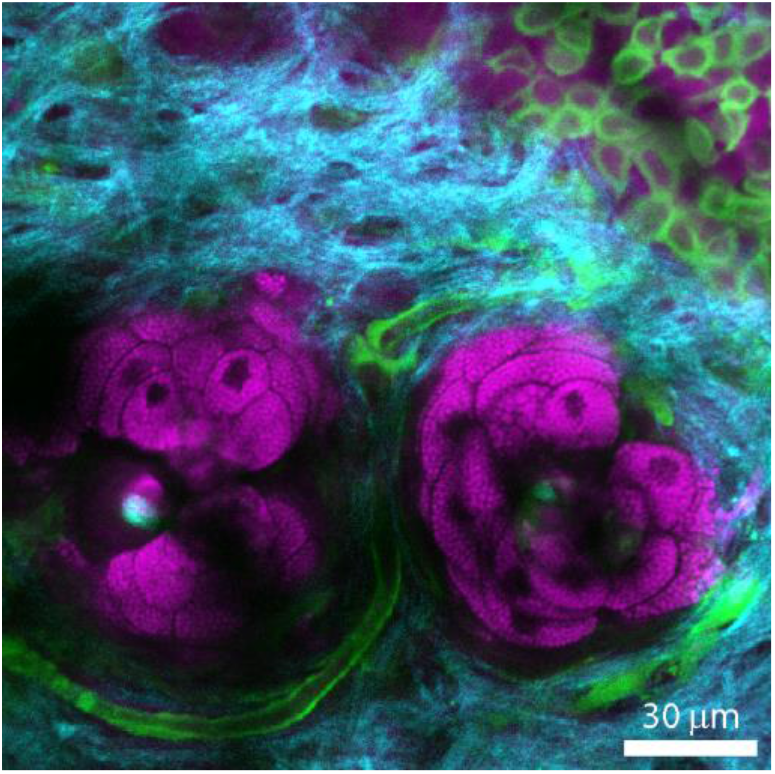
Combined SHG (cyan), GCaMP6s fluorescence (green) and epi-CARS (magenta) *in vivo* imaging of mouse earlobe skin showing, respectively, collagen, keratinocytes plus blood vessels, and sebocytes of sebaceous glands. λ_1_ = 840 nm, P_1_ = 12 mW, λ_2_ =1105 nm, P_2_ = 15 mW. The SHG signal was collected around 420 nm; the GCaMP6s 2c-2PE fluorescence signal was collected from 505 nm to 545 nm; the CARS signal was collected around 678 nm; pixel size=192 nm.

**Fig. 9.**
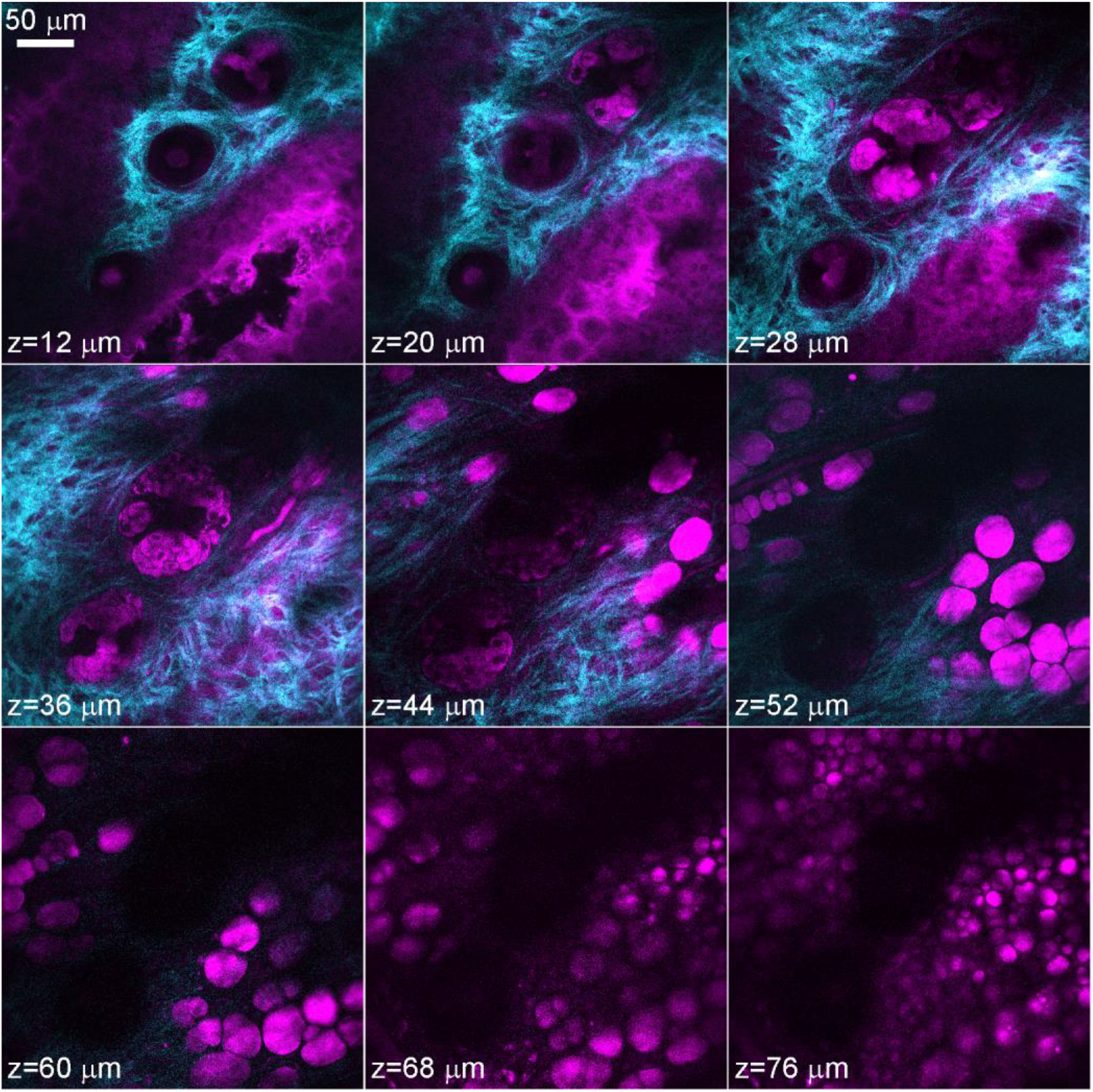
Through-focus image sequence combining SHG (cyan) and epi-CARS (magenta) *in vivo* signals of unstained wild type mouse earlobe skin showing collagen fibers, keratinocytes, sebocytes of sebaceous glands, blood vessels and lipid droplets of the adipose tissue. Pixel size = 309 nm; other parameters as in Fig. 8.

CARS imaging highlighted also dermal vasculature and flowing erythrocytes (Figure 10), suggesting that this label-free imaging modality can be used to quantitatively evaluate blood flow at the level of individual capillaries.

**Fig. 10.**
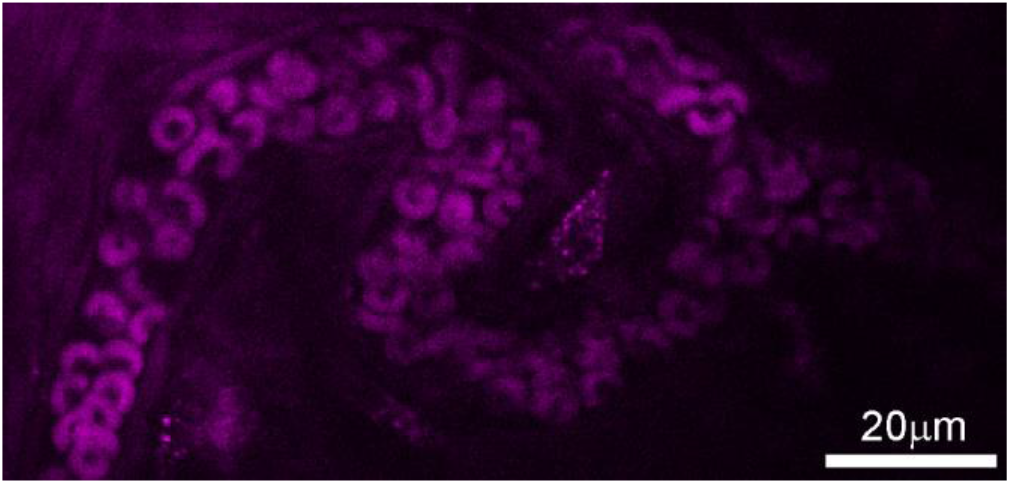
CARS imaging of flowing erythrocytes in live mouse capillary. The image was acquired using λ_1_ = 840 nm, λ_2_ =1105 nm, P_1_ = 12 mW, P_2_ = 22 mW. The CARS signal was acquired around 678 nm; pixel size=219 nm. The image was obtained by averaging 11 consecutive video frames.

## 4. Conclusions

The results presented above demonstrate that a relatively modest investment in terms of additional equipment can substantially improve and expand the performances of a commercial multiphoton / STED architecture. The key points that allowed us to meet our goals were: i) the addition of the delay line with computer-controlled sub-micrometer precision and related optics to compensate for the 5.5 ns delay between Ti:Sa and OPO pulses and Ti:Sa beam divergence; ii) the insertion of high-precision remotely-controlled beam-stirring mirrors for accurate spatial alignment of Ti:Sa, OPO and STED beams; iii) the customized feature of the control software that allowed us to move the vortex phase plate in two orthogonal directions within the commercial STED module; iv) the picosecond delayer electronics for accurate timing of the STED pulses relative to the Ti:Sa clock. The added benefits include not only the possibility to perform 2PE-STED microscopy up to a depth of 70 μm, but also 2c-2PEF simultaneously with label-free imaging deep in tissues to visualize structures and fast dynamic processes *in vivo*.

## Acknowledgments

FM was the recipient of a Shanghai Thousand Talent Honor from the Shanghai Municipal Government.

